# *PIK3C2B* promotes epithelial to mesenchymal transition and EGFR inhibitors insensitivity in epidermal squamous cell carcinoma

**DOI:** 10.1101/363721

**Authors:** Silvia Crespo Pomar, Anna Borgström, Alexandre Arcaro, Roch-Philippe Charles

## Abstract

While the class I of PI3Ks has been deeply studied due to its clear implication in cancer development, little is known about the class II of PI3Ks. However, recent accumulation of data is now revealing that PI3KC2β, one isoform of this class of PI3Ks, may also play a role in cancer. Specifically, recent studies have suggested an implication of PI3KC2β in metastasis formation through the promotion of epithelial to mesenchymal transition (EMT). Here, we report that the overexpression of PI3KC2β in the epidermal squamous cell carcinoma (ESCC) cells A431 promotes apparent EMT transformation. We further confirm this EMT by showing modification in several biochemical markers (E-cadherin, β-catenin, Snail, Twist1 and Vimentin). Furthermore, an intracellular co-localization of E-cadherin, β-catenin and EGFR was observed. This transformation decreased EGFR signaling and the sensitivity to inhibitors targeting this receptor. To confirm our results, we have used the colon adenocarcinoma cells HT29 and induced overexpression of PI3KC2β in these cells. We could recapitulate in this model some of our major findings regarding EMT in the PI3KC2β overexpressing A431 cells. Taken together, these data support a role of PI3KC2β in promoting EMT.

## Introduction

Phosphoinositide-3-kinases (PI3Ks) are a family of lipid kinases that play a role in coordinating main intracellular signaling pathways following the activation by upstream agonists such as receptor tyrosine kinases (RTKs) or G protein coupled receptors (GPCR) (1). They phosphorylate the 3’-hydroxyl group of the inositol ring of three species of phosphatidylinositol (PI) lipid substrates. This triggers the formation of the second messengers PI(3)P (phosphatidylinositol 3-phosphate), PI(3,4)P2 (phosphatidylinositol 3,4-bisphosphate) and PIP3 (phosphatidylinositol 3,4,5-trisphosphate), specifically enriched in different cellular compartments (1). Subsequently, different effectors will be recruited and bind to these PI3K phospholipid products, changing their conformation and inducing the propagation of signals involved in cell cycle progression, cell growth, survival, migration, and intracellular vesicular transport (2).

In mammals, the PI3K family is formed by eight different catalytic PI3K isoforms, classified into three classes (I, II and III), based on sequence homology and *in vitro* substrate specificity (2). The different PI3K isoforms can be expressed in a tissue-specific manner and seem to have non-redundant roles. Among the PI3K classes, mutations in class I PI3K isoforms are especially linked with cancer development, being *PIK3CA* the second most frequently mutated oncogene (3). However, while class I PI3Ks have been well studied, little is known about the function of class II. This is mainly because they were first identified by PCR and homology cloning approaches, not based on their cellular function (4,5). Finally, the lack of specific inhibitors and mouse models hindered the discrimination between the biological functions of each isoform.

PI3KC2β is one of the three isoforms from class II PI3K. It is ubiquitously transcribed, showing the highest levels in placenta and thymus (6,7). This lipid kinase is a monomer mainly activated downstream of RTKs, like EGFR, and other GPCRs (8,9). It has a substrate specificity directed towards PI and PI(4)P, generating a pool of PI(3)P or PI(3,4)P_2_ in the plasma membrane and endosomes (10,11). Through the recruitment of different secondary messengers and the subsequent activation of AKT/mTOR or RAC signaling pathways, PI3KC2β regulates different cellular functions like cell migration, cell growth, cell survival, invasion, cell cycle progression and K^+^ channel activation (12–15).

PI3KC2β is the isoform of the class II of PI3Ks that has the most documented implication in cancer development. Its overexpression at the protein and mRNA level in a variety of human tumors (12,13,16,17), and the promising effects of PI3KC2β inhibition observed in leukemia, brain and neuroendocrine tumors (18), are a few of the observations that support this hypothesis. Recent studies have observed a correlation between PI3KC2β expression and metastasis formation in breast, prostate cancer and esophageal squamous cell carcinoma (12,13,19).

Most of cancer-related death are due to metastases formation. To that end, primary tumors need to undergo the biological process known as epithelial-mesenchymal transition (EMT). This process allows immobile epithelial cells to acquire a mobile mesenchymal phenotype, losing cell-cell contact dependency. This transformation is promoted by the expression of several transcription factors like Snail, Slug, and Twist. Two important mechanisms are the replacement of keratin cytoskeleton by a more plastic Vimentin cytoskeleton and the decreased membranal expression of cell–cell adhesion proteins such as E-cadherin or β-Catenin (20). Different studies have suggested an implication of PI3KC2β in EMT (12,13). However, the exact mechanism by which this kinase is participating in this process is still unknown.

In the epidermal squamous cell carcinoma (ESCC) cells A431, the overexpression of PI3KC2β has been reported to enhance membrane ruffling, migration speed of the cells, protection from anoikis and cell proliferation (14). We decided to assess if some of these previously reported effects could be consequence of a possible process of EMT induced in these cells after the overexpression of PI3KC2β. For this purpose, A431 cells stably expressing PI3KC2β were evaluated for different EMT markers expression. The cytoplasmic localization of different cell-cell adhesion proteins was also visualized. Moreover, the functional impact of PI3KC2β overexpression was assessed in relation to its effect in EGFR signaling and the sensitivity to inhibitors targeting this receptor. Finally, to further validate our results, PI3KC2β overexpression was induced in an additional cell line, HT29.

## Materials and methods

### Cell Lines

A431 human epidermoid carcinoma cells and HT29 cells were purchased from the American Type Culture Collection. A431 cells were grown in DMEM (Sigma Aldrich, cat. no. D5796) supplemented with 10% (v/v) fetal bovine serum (FBS) (Gibco, cat. no. 10082147), 2 mM L-glutamine (Gibco, cat. no. 25030081) and 50.000 units of penicillin/streptomycin (Gibco, cat. no. 15140122). HT29 cells were grown in McCoy’s 5A Medium (Gibco, cat. no. 26600-023) supplemented with 10% (v/v) fetal bovine serum (FBS) (Gibco, cat. no. 10082147), 2 mM L-glutamine (Gibco, cat. no. 25030081) and 50.000 units of penicillin/streptomycin (Gibco, cat. no. 15140122). Cells were passaged every 3 to 4 days and were kept incubated in a humidified atmosphere of 5% CO_2_ at 37 °C. Stably transfected HT29 and A431 clones were grown in the presence of 0.8 mg/ml G418 (Life Technologies).

### Stable transfection

A431 cell lines stably expressing PI3KC2β were generated as previously described (8). HT29 cell lines stably expressing PI3KC2β were generated as follows. The cDNA of N-terminal myc-tagged (MEQKLISEEDL) *PI3KC2β* wild-type was cloned into pcDNA3 vector (Invitrogen) using EcoRI and XhoI sites as described in Arcaro et al. 1998. The transfection of the pcDNA3-PI3KC2β plasmid was performed using Lipofectamine 2000 (Invitrogen) according to manufacturer’s instructions. 48 h post-transfection cells were split into selection medium containing 1 mg/ml G418. Cells were cultured in the selection medium for 2-3 following weeks. Medium was changed each 72 h. When the single G418 resistant colonies appeared, they were further selected and expanded. After 2-3 passages, expression of PI3KC2β was verified by qPCR and western blot.

### Western blotting

Proteins were extracted in RIPA buffer (20 mM Tris-base pH=8, 150 mM NaCl, 1% TritonX-100, 0.1% SDS, 0.5% sodium deoxycholate. Sigma, cat. no. D6750-10G) supplemented with Halt™ protease/phosphatase inhibitor cocktail (Pierce, cat. no. 78444). Protein concentration was assessed by Pierce BCA protein assay kit (Thermo Scientific, cat. no. 23225) and 20 μg of total proteins were
separated by SDS-PAGE. Protein gels were transferred into nitrocellulose membranes and blocked with Tris buffered saline (TBS 1x 130 mM, NaCl 30 mM, Tris-Cl pH=7.5) containing 5% Bovine Serum Albumin (BSA) for 2 h. Western blots were probed with rabbit anti-PI3KC2β polyclonal antibody (1/1000, described in (6), rabbit anti-pAKT S473 (1/1000, Cell signaling, cat. no. 4090), rabbit anti-pERK (1/2000, Cell signaling, cat. no. 4370), mouse anti-panAKT (1/1000, cat. no. 2920, Cell signaling), mouse anti-ERK (1/5000, Cell signaling, cat. no. 9107), mouse anti-β-ACTIN antibody (1/15000, Sigma-Aldrich, cat. no. A5316), mouse anti-E-cadherin (1/1000, Abcam, cat. no. 76055), rabbit anti-β-catenin (1/1000, Abcam, cat. no. 32572) and rabbit anti-EGFR (1/1000, Cell signaling, cat. no. 4267).

Primary antibodies were detected using goat anti-rabbit IR680 (1/10’000, Li-Cor Bioscience, cat. no. 926-68071) and goat anti-mouse IR800 (1/10’000, Li-Cor Bioscience, cat. no. 926-32210) and imaged by a LI-COR OdysseySa^®^ scanner. Antibodies were diluted in TBS x1 containing 2% BSA, 0.1% Tween and 0.1% sodium azide.

### Proliferation assays

The EGFR kinase inhibitors, Gefitinib and Erlotinib, were purchased from ChemieTek (Indianapolis, IN, USA). Cells were seeded at a density of 3-5 × 10^3^ cells/well (depending on the cell line) in a 96-well plate using the cell line-specific culture medium. Cells were allowed to adhere overnight and were then treated with the indicated concentration of drugs for 72 h. After the treatment, cells were fixed with 10% buffered formalin, stained with 0.2% crystal violet (Sigma-Aldrich, cat. no. C3886-25G) in 2% ethanol, washed 5 times in dH_2_O and lysed with 100 μL 1% SDS to recover the dye. Optical density (OD) at 550 nm was measured with a microplate reader and normalized to the DMSO control. Dose-response curves were carried out by using the medium described previously.

### Quantitative Real-time PCR

RNA was extracted using the RNeasy Mini Kit (Qiagen, cat. no. 74106) according to the manufacturer’s protocol. Reverse-transcription was performed with Superscript II Reverse Transcriptase (Invitrogen, cat. no. 18064-014) following the manufacturer’s protocol. The Sybr green^®^ real-time primers used in this project were purchased from Applied Biosystems and are depicted in Table 1. PCR reactions were performed in a ViiA7 cycler (Applied Biosciences) using SybrSelect Mastermix (Applied Biosystems, cat. no. 4472908). Expression of mRNA was normalized to *ACTB* and *GAPDH* housekeeping genes using the 2^ΔΔct^ method.

**Table 1:**
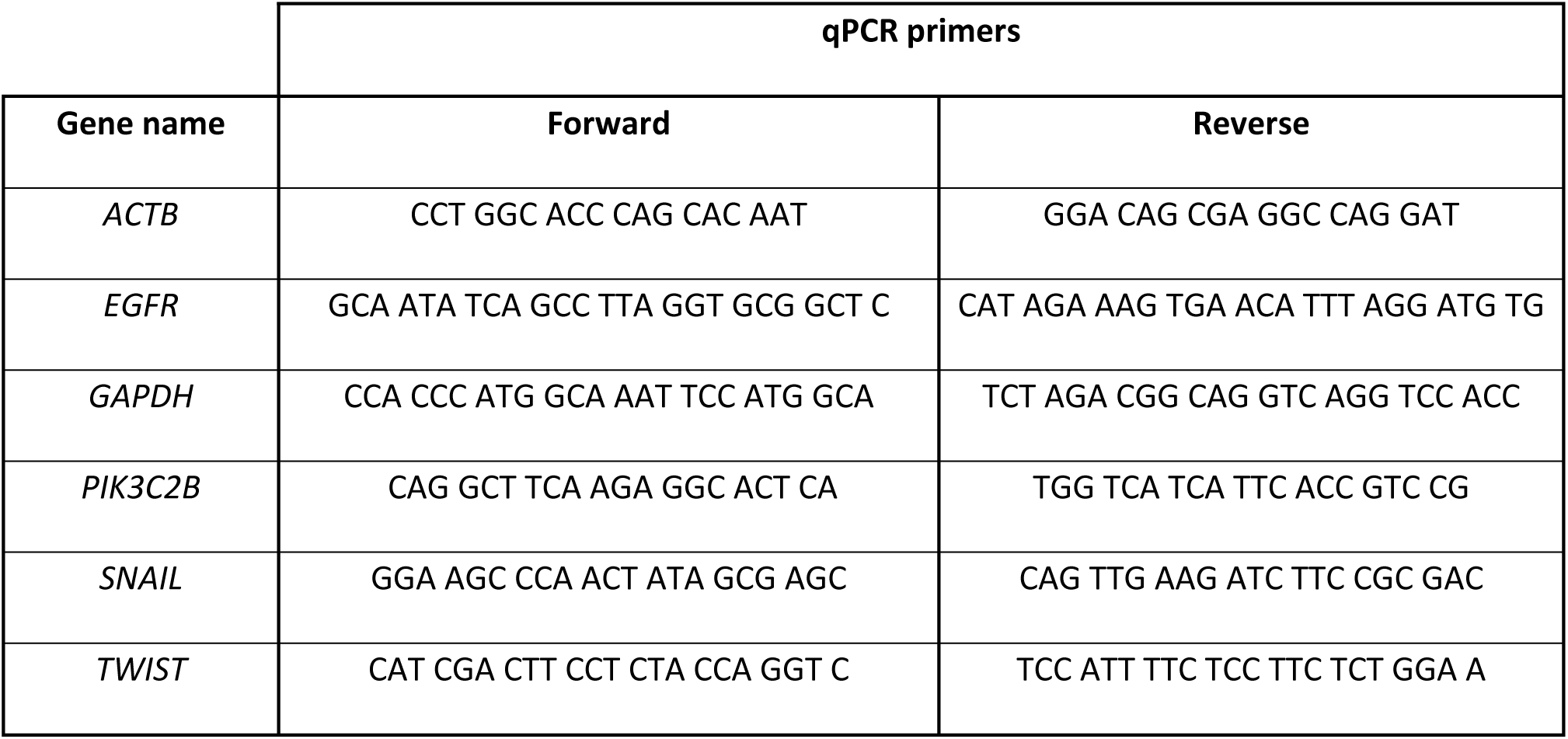
list of primers used for qPCR quantifications.

### Immunofluorescence staining

Cells were grown for 24 h on glass coverslips in 24 well plates to around 60% confluence. After 10% formalin (Sigma Aldrich, cat. no. HT501128) fixation (10 min), coverslips were washed 3×10 min in phosphate buffered saline (1x PBS: 137 mM NaCl, 2.7 mM KCl, 18 mM KH_2_PO_4_, 10 mM Na_2_HPO_4_) and cells were subsequently permeabilized with a 1x PBS, 0.3% TritonX-100 solution. Following blocking with a 1% BSA, 0.2% gelatin, 0.05% saponin in 1x PBS solution and washing with a 0.1% BSA, 0.2% gelatin, 0.05% saponin in 1x PBS solution. Cells were incubated overnight at 4 °C with the primary antibodies, rabbit anti-Vimentin (1/300, Abcam, cat. no. 5741,), mouse anti-E-cadherin (1/200, Abcam, cat. no. 76055), rabbit anti-β-catenin (1/200, Abcam, cat. no. 32572), rabbit anti-EGFR (1/300, Cell Signalling cat. no. 4267), rabbit anti-PI3KC2β polyclonal antibody (1/200, described in (6)), mouse anti-58K (5 μg/ml, cat. Abcam, no. 27043,), mouse anti-RAB 7 (1/300, cat. no. 376362, Santa Cruz Biotechnology) and mouse anti-RAB 11 (1/300, Cell Signalling, cat. no. 5589). The secondary antibodies, goat anti-rabbit Alexa-488 (1:400, Life Technologies, cat. no. A11034) and goat anti-mouse Alexa-633 (1:300, Life Technologies, cat. no. A21052,) were used to detect antigen-antibody complexes; slides were counter-stained with DAPI (500 ng/ml, Sigma Aldrich, cat. no. 32670-25MG) to visualize DNA.

### Slide Scanning

Slides were scanned using a Panoramic Midi digital slide scanner (3DHISTECH) and analyzed with the Panoramic Viewer software (3DHISTECH). For the quantification of Vimentin/DAPI positive cells, an area of the slide was selected and analyzed with CellQuant software (3DHISTECH).

### Confocal images

Cells were examined by confocal Leica SP8 X STED inverted microscope under 63× magnifications. Image files were collected as a matrix of 1024 × 1024 pixels and analyzed using the Leica Application Suite X Microscope Software.

### Wound healing assay

Cells were grown to confluency in 6-well tissue culture plates. The resulting cell monolayer was scratched with a 200 μL pipette tip generating two parallel wounds and returned to a humidified incubator at 37 °C 5% CO_2_. Two replicates for each treatment and four phase-contrast images were acquired along the length of each wound every 2-4 h under 40x magnifications. Cell migration was also assessed in a 96-well format using ORIS cell migration assay from Platypus Technologies (cat. no. CMA1.101). Cells were seeded at a density of 7×10^4^ cells/well in 100 μl of medium around the stoppers and left to attach for 24 h. Next day, the stoppers were removed, and phase-contrast images were acquired from each well every 2-4 h. The wound area was determined from these images after delineating the edges digitally using Adobe Photoshop CS4 software.

### Statistical analysis

Data is presented as average ± SD. All experiments were performed in triplicates. Statistical analyses were conducted using GraphPad Prism 7 (GraphPad Software). The statistical test used is indicated in the respective figure legend. P < 0.05 was considered as statistically significant.

## Results

### The overexpression of PI3KC2β changes the migration front of A431

To further study the implication of PI3KC2β in regulating cell migration (14,21), stably transfected A431 cells were used. First, *PIK3C2B* overexpression was validated at the protein level. Results of the conducted western blots indicated strong PI3KC2β overexpression in stably expressing clones, increasing the levels of this kinase to 233.88 ±16.95% of the parental cell line expression level (Fig 1A). We then tested the effect of this overexpression in terms of cellular migration. The PI3KC2β overexpressing cells presented increase in wound healing closure speed compared to the parental cell line A431 (Fig 1B). During these migration assays, A431 cells appeared to have tighter cell-to-cell junctions, typical from epithelial cells, whereas A431C2β showed a more fibroblastic spindle-like morphology (Fig 1C).

**Figure 1:**
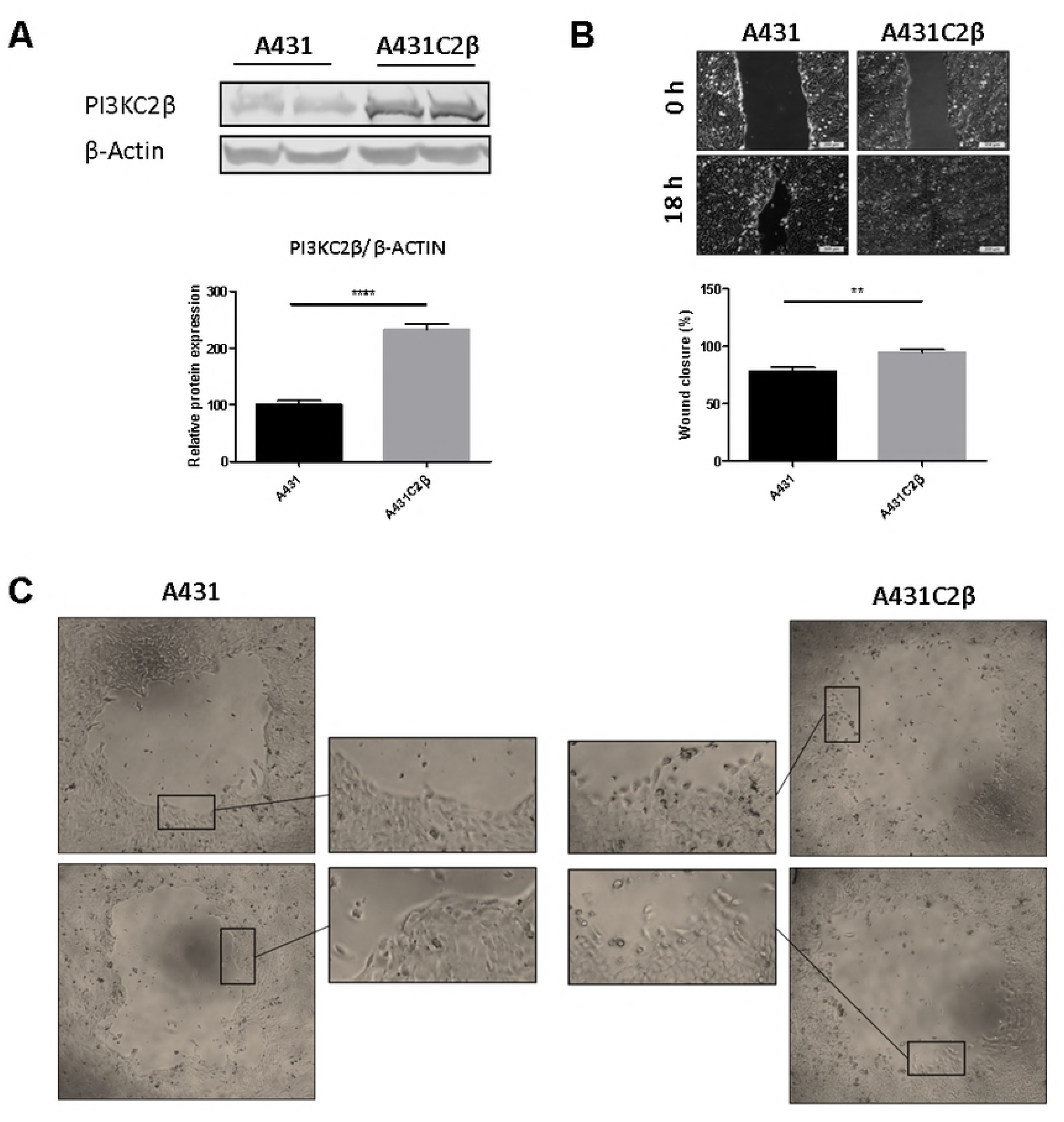
The overexpression of PI3KC2β changes the migration front of A431. **A** Western blot and Relative protein expression of PI3KC2β in A431 and A431C2β cells. **B** Wound healing closure in A431 and A431C2β cells after 18 h of generating a wound with a 200 μL pipette tip. Images are representative of 4 different experiments performed in a 6 well plate. **C** Comparison of the cell morphology during cell migration of A431 and A431C2β cells in 96 well plates. Representative images obtained through phase-contrast microscopy. Means ±SD; n = 4 independent experiments; unpaired two tailed t-test.

### PI3KC2β overexpression increases EMT markers expression

Following the primary observation, the levels of EMT markers in A431 and A431C2β cells were evaluated. E-cadherin and β-catenin expression were reduced in A431C2β cells compared to A431 by 21% and 71% respectively (Fig 2A). Snail and Twist, two transcription factors involved in EMT presented elevated transcription with almost 4 folds for *SNAI1* and more than 3 folds for *TWIST1* (Fig 2B) of the parental transcription levels in the A431C2β. Finally, immunostaining of Vimentin (green) showed a higher positivity index (more than 7 folds higher) in A431C2β compared to A431 (fig 2C).

**Figure 2:**
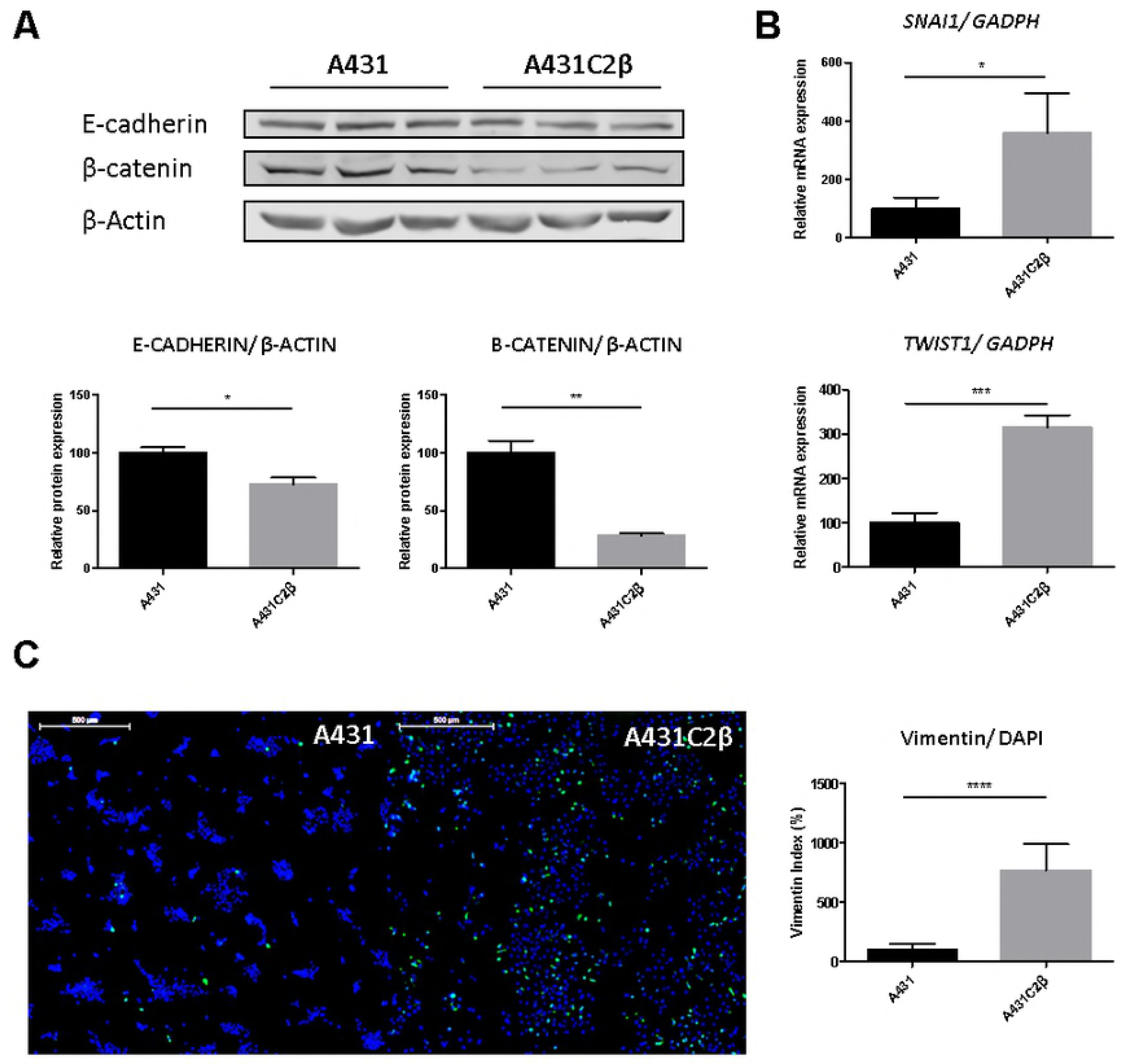
The overexpression of PI3KC2β increases the expression of EMT markers in A431. **A** Western blot and relative protein expression of E-cadherin and β-catenin in A431 and A431C2β cells. **B** Relative mRNA expression of *SNAI1* and *TWIST1* in A431 and A431C2β cells. **C** Expression of Vimentin visualized by immunofluorescence in A431 and A431C2β cells. Nuclear staining with DAPI (blue) and anti-Vimentin antibody (green). The results of A431C2β were normalized to A431 samples. Means ±SD; n ≥ 3 independent experiments; unpaired two tailed t-test. Size bar = 500μm.

### Cytoplasmic co-localization of E-Cadherin, β-Catenin in A431C2β

To study further the effect of PI3KC2β overexpression, the cellular localizations of E-cadherin and β-catenin were visualized by confocal microscopy after immunofluorescence staining. While in A431 cells, E-cadherin and β-catenin showed staining mainly at the plasma membrane, A431C2β cells, showed staining at both the plasma membrane and in a cytoplasmic region close to the nucleus (Fig 3A).

**Figure 3:**
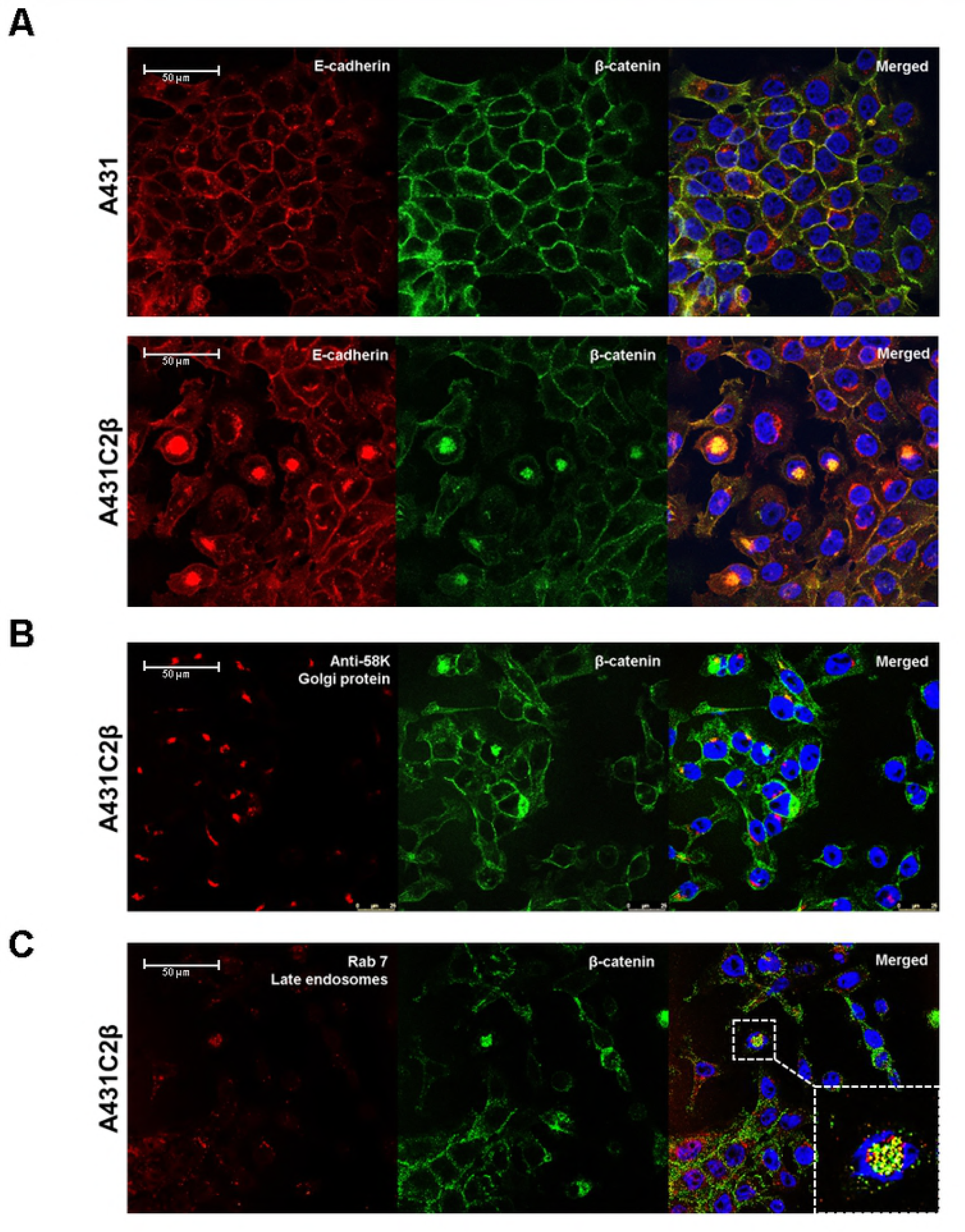
Cytoplasmic co-localization of E-cadherin, β-catenin in A431C2β. Dual-color immunofluorescence co-localization was performed in A431 and A431C2β cells grown on glass coverslips. Images were obtained by confocal microscopy. **A** Expression of E-cadherin (red) and β-catenin (green) visualized with immunofluorescence in A431 and A431C2β cells. **B** Expression of 58K, a golgi marker (red) and β-catenin (green) visualized with immunofluorescence in A431 and A431C2β cells. **C** Expression of Rab7, a late endosome marker (red) and β-catenin (green) visualized with immunofluorescence in A431 and A431C2β cells. Nuclei counterstain with DAPI (blue). Size bar = 50μm.

To better define the cytosolic sub compartment of the cell where E-cadherin and β-catenin were localized, dual-color immunofluorescence stainings of β-catenin with either 58K (22) used as a marker of the Golgi apparatus (Fig 3B) or Rab 7 (23) as a marker for late endosomes (Fig 3C) were performed. β-catenin co-localized with neither 58K nor Rab 7.

### Cytoplasmic co-localization of PI3KC2β and EGFR with E-cadherin/β-catenin aggregates

In attempt to explain the effect of PI3KC2β-induced cytoplasmic localization of E-cadherin and β-catenin, we tested if PI3KC2β localization was overlapping with E-cadherin and β-catenin. Dual-colour immunofluorescence staining of PI3KC2β with E-Cadherin was also performed revealed partial co-localization of these proteins in the cytoplasm (Fig 4A).

**Figure 4:**
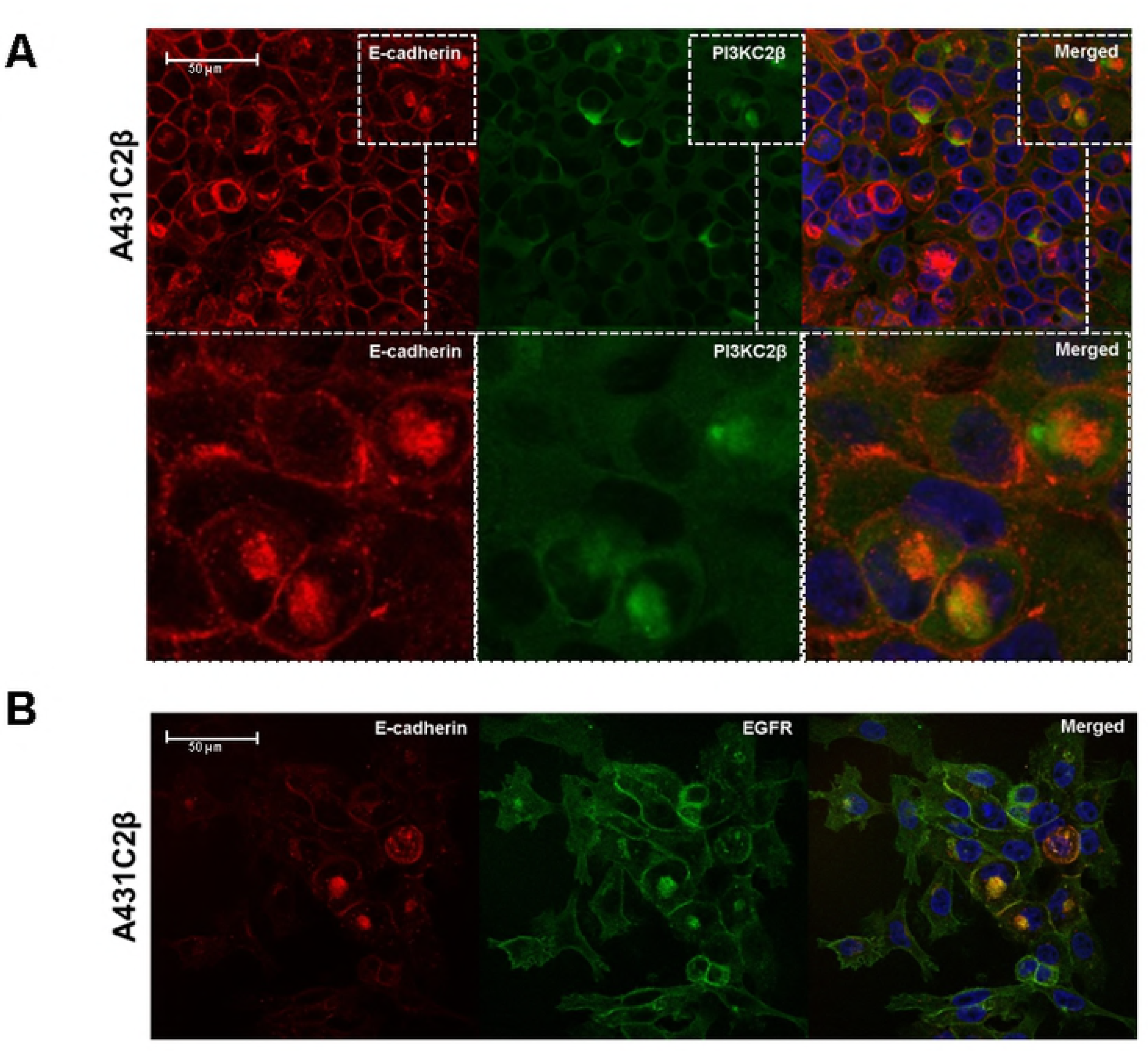
Cytoplasmic co-localization of PI3KC2β and EGFR with E-cadherin/β-catenin aggregates. Dual-color immunofluorescence co-localization was performed in A431 and A431C2β cells grown on glass coverslips. Images were obtained by confocal microscopy. **A** Expression of E-cadherin (red) and PI3KC2β (green) visualized with immunofluorescence in A431C2β cells. **B** Expression of E-cadherin (red) and EGFR (green) visualized with immunofluorescence in A431C2β cells. Nuclei counterstain with DAPI (blue). Size bar = 50μm.

We performed here a dual-color immunofluorescence staining of EGFR with E-cadherin (Fig 4B). The overlap of the two signals in the cytoplasm showed that the two proteins co-localize suggesting that PI3KC2β overexpression is leading to the internalization of E-cadherin, β-catenin and EGFR together in the same intracellular compartment.

### PI3KC2β overexpression reduces EGFR signaling

The PI3KC2β-driven relocalization of EGFR has been further examined. Quantitative PCR analysis showed a significant reduction of *EGFR* transcription (−88.47 ±18.87%) but also a significant reduction of EGFR expression (−81.38 ±7.42%) in the A431C2β cells (Fig 5A and 5B) compared to the parental cell line A431. Consequently, alterations in EGFR downstream signaling were analyzed. We could find that A431C2β cells showed decreased phosphorylation levels for AKT (29.81 ±0.69%) and ERK (22.68 ±4.72%) (Fig 5C) compared to their respective total expression. This reduction was less remarkable when compared to beta-Actin, since a strong increase of total-ERK was also found (Fig 5C).

**Figure 5:**
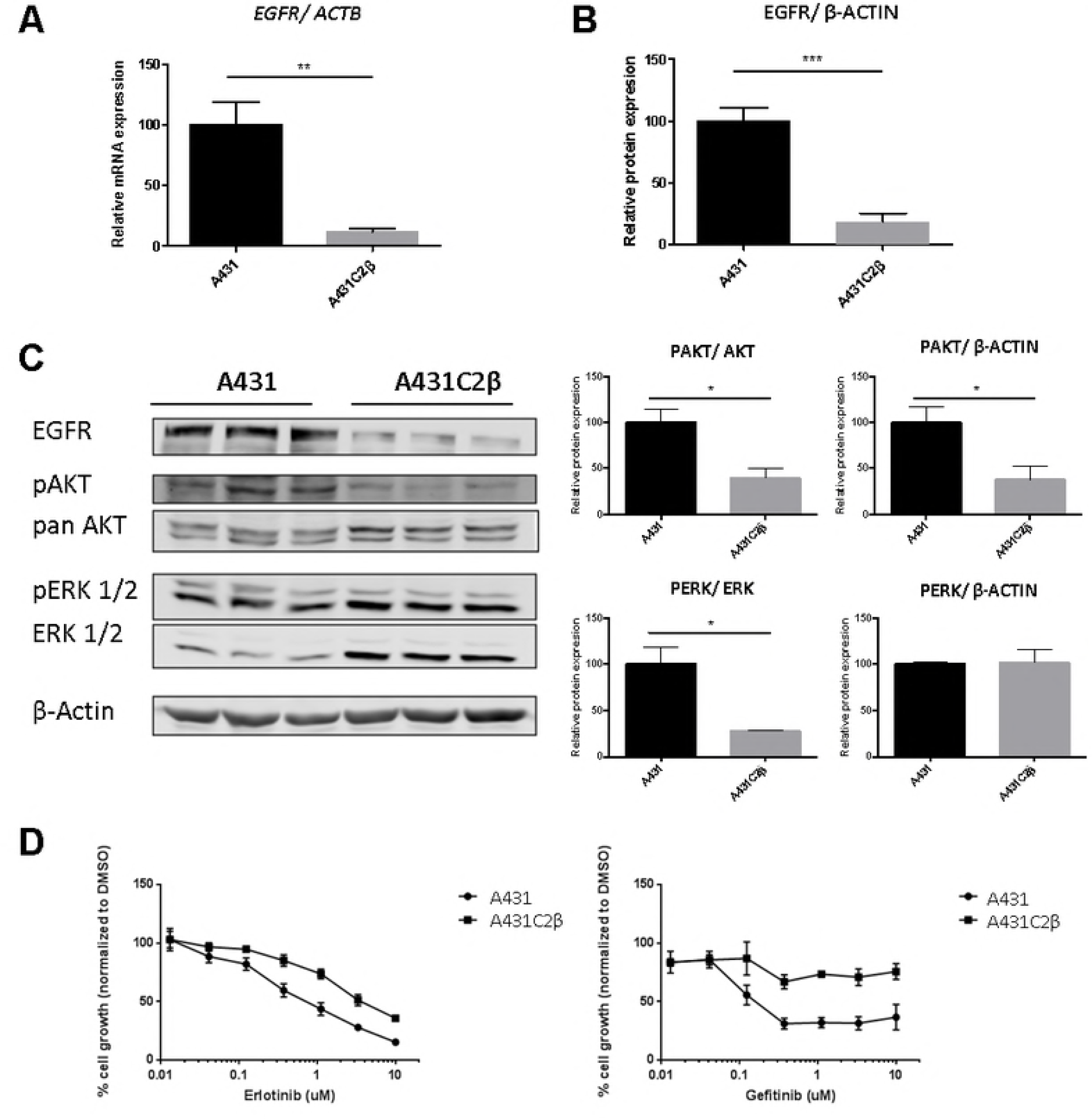
The effect of PI3KC2β overexpression in EGFR signalling. *EGFR* transcription (**A**) and EGFR expression (**B**) compared by qPCR and western blots in A431C2β cell line compared to parental cell line. **C** Western blot picture and analyses for EGFR, AKT phosphorylation and ERK phosphorylation in A431 and in A431C2β. **D** A431 and A431C2β, dose response to EGFR TKIs, Erlotinib and Gefitinib. Cell viability was measured via Crystal violet assay. Means ±SD; n = 3 independent experiments; unpaired two tailed t-test.

PI3KC2β has also been suggested to participate in chemo-resistance to different drugs (6–8). We therefore tested if this PI3KC2β-driven relocalization of EGFR translates into modified response to EGFR kinase inhibitors. Cell proliferation of the two lines, A431 and A431C2β, was measured after 72 h of treatment with 0.1% DMSO or with increasing concentrations of EGFR kinase inhibitors, Erlotinib and Gefitinib. Results obtained from the crystal violet assay showed that PI3KC2β overexpression increases Erlotinib and Gefitinib resistance (Fig 5D). More specifically, the measured IC50 for Erlotinib shifted from 0.652 μM to 4.434 μM between A431 and A431C2β, and the measured IC50 for Gefitinib shifted from 0.084 μM to 0.47 μM.

### The overexpression of PI3KC2β induces also EMT markers expression in HT29 cells

To confirm our observations, HT29 cells were stably transfected with the same Myc-tagged *PIK3C2B* expression vector used for the A431 cells. After transfection, the different generated clones were evaluated on a transcriptional level for *PIK3C2B* overexpression. The clone HT29C2β (1/14) was selected for its highest expression of *PIK3C2B* with a relative expression of 751 ±81.96% in comparisons to the parental HT29 cell line (Fig 7A) or the cells transfected with the empty pcDNA3 vector. The levels of different EMT markers were also evaluated by western blot. In HT29 cells *PIK3C2B* overexpression resulted in a decrease in the expression levels of E-cadherin (−40.05 ±8.62%) in relation to the parental HT29 cell line (Fig 6A). Furthermore, HT29C2β cells also showed increased mRNA levels for *SNAI1* (2.5 folds) (Fig 6B). Finally, the comparison of the morphology of the migration front between HT29 and HT29C2β cells showed in HT29 an apparent tighter cell-to-cell contacts than in HT29C2β (Fig 6C).

**Figure 6:**
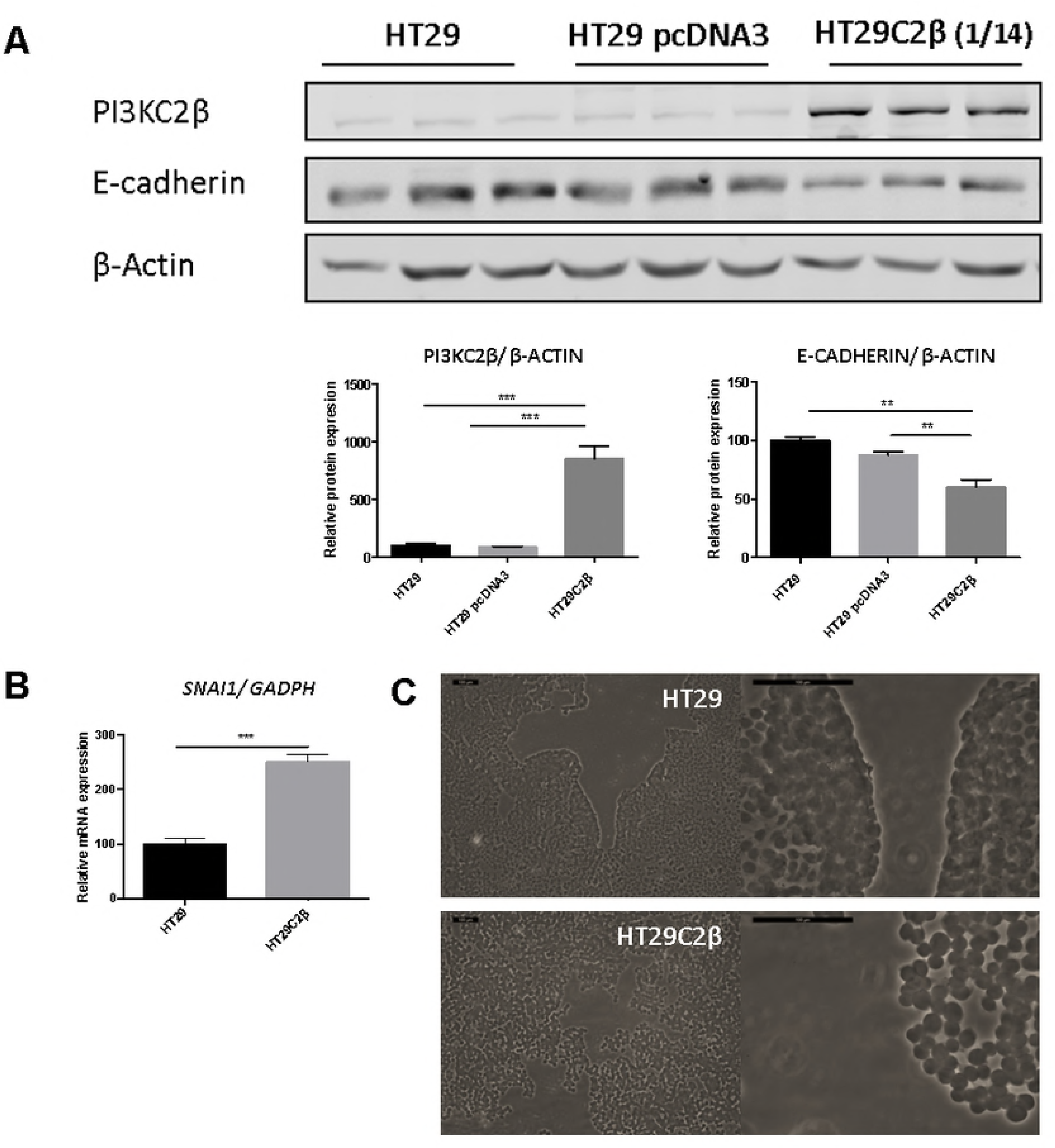
The overexpression of PI3KC2β induces the expression of some EMT markers in HT29. **A** Relative protein expression of E-cadherin in HT29 cells, HT29 cells transfected with pcDNA3 plasmid, and HT29C2β (clone 1/14) cells transfected with a pcDNA3-PI3KC2β plasmid (evaluated by Western blot). **B** Relative mRNA transcription of *SNAI1* in HT29 and HT29C2β cells (evaluated by qPCR). The results were normalized to HT29 samples. Means ± SD; n ≥ 3 independent experiments; unpaired two tailed t-test. **C** Comparison of the cell morphology during cell migration of HT29 and HT29C2β. Representative images obtained through phase-contrast microscopy.

**Figure 7:**
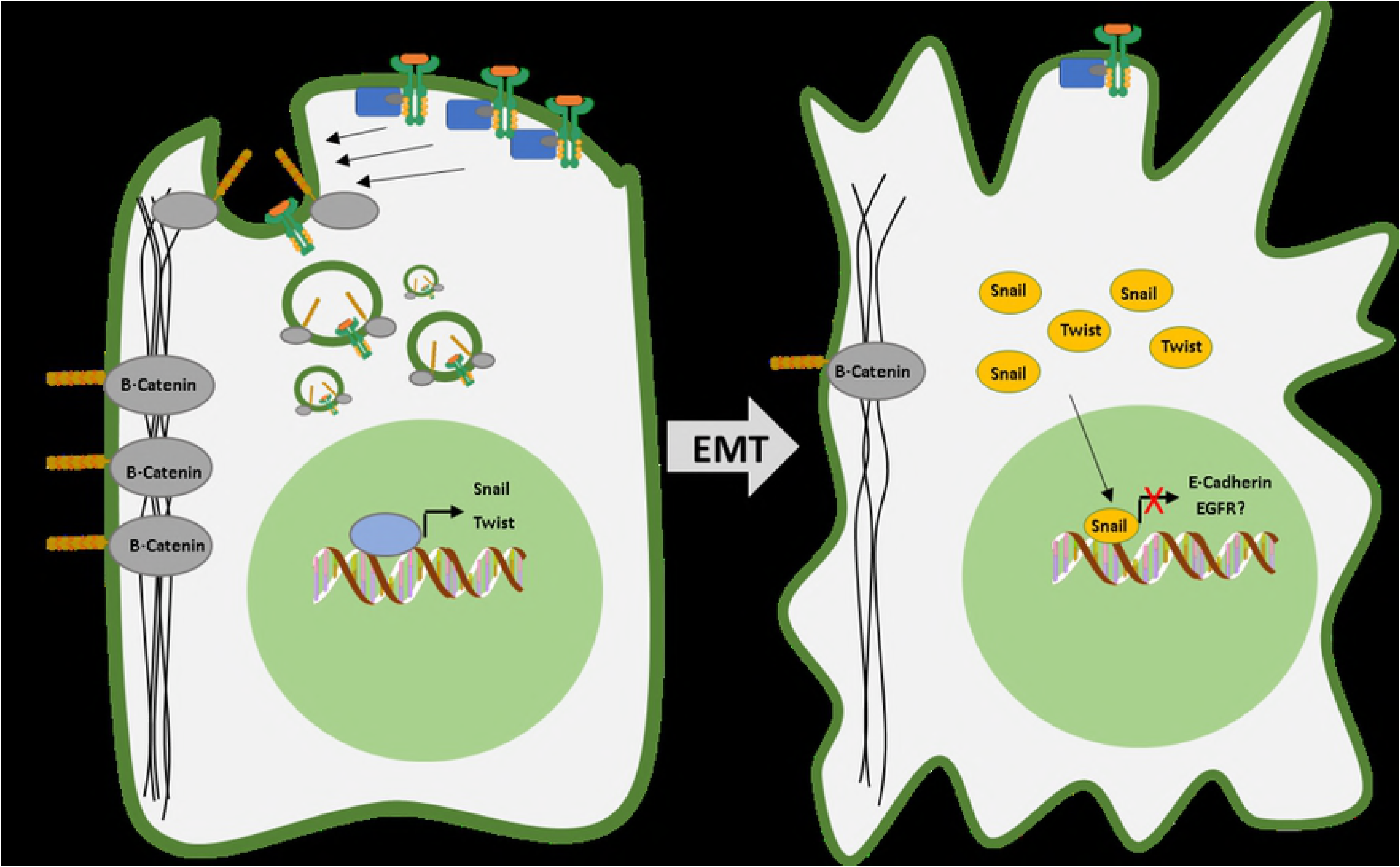
Proposed model of how the overexpression of PI3KC2β affects intracellular vesicular trafficking of E-cadherin, β-catenin and EGFR and promotes EMT. <u>Left:</u> Early phase of EMT: PI3KC2β induces the endocytic internalization of EGFR, E-cadherin and β-catenin removing these adherent components and the receptor from cell surface. Additionally, the overexpression of PI3KC2β contribute to the activation of the expression of different transcription factors like Twist and Snail. This constitutes the transcriptional machinery that will lead to EMT. <u>Right:</u> Late phase of EMT: Twist and Snail will repress the expression of E-cadherin and EGFR. The reduced expression of E-cadherin destabilizes cell-cell adhesions promoting a change in the morphology of the cell.

## Discussion

In this study, we have confirmed that PI3KC2β overexpression in A431 epidermoid carcinoma cells is promoting a more motile phenotype increasing wound healing closured (Fig 1B). Katso *et al*. previously reported that the increased expression of PI3KC2β stimulates Rac activity in these cancer cells, increasing membrane ruffling and migration speed of the cells (14). Moreover, the overexpression of this PI3KC2β also renders A431 cells resistant to anoikis (14). Additionally, in the present study we have observed a change in the morphology of the cells from the migration front, that switch from apparently tight cell-to-cell contact to a more fibroblastic spindle-like morphology after PI3KC2β overexpression (Fig 1C). This morphology change resembles the cytoskeleton reorganization typical of a EMT process (24,25).

We observed that the overexpression of PI3KC2β induced an increased expression of different EMT markers: mRNA levels of Snail and Twist, increase positivity index of Vimentin and decreased protein levels of E-cadherin and β-catenin (Fig 2). This is consistent with what has been already observed, in prostate cancer that PI3KC2β has been reported to control cell invasion by regulating SLUG expression, a transcription factor promoting EMT progression (13). Additionally, in breast cancer cell lines, PI3KC2β regulates cell invasion and activates the transcription factor STAT3, that controls the expression of master EMT transcription factors (12).

In A431, the induction of EMT can be elicited by a chronic EGF treatment that promotes the endocytosis of E-cadherin followed by the dissociation of the E-cadherin/β-catenin complex and the subsequent trans-activation of the β-catenin/lymphoid enhancer factor 1 (LEF-1) pathway (26,27). We are reporting here that PI3KC2β overexpression is promoting a perinuclear localization of E-cadherin, β-catenin and EGFR instead of mainly at the plasma membrane (Fig 3A, 4A). However, we were not able to determinate their specific cytoplasmic localization as they do not co-localize with markers of Golgi apparatus or late endosomes (Fig 3B-C). Despite this, the proximity to late endosomes markers suggest that these molecules could be localized in another type of endosomes. A similar phenotype was observed in A431 after the treatment with lysophosphatidic acid (LPA) (28). In this study E-cadherin and β-catenin were colocalized by immunostaining in a discrete region near the nucleus that was identical to the perinuclear endocytic recycling compartment (ERC). The authors proposed that the ERC may be a site of residence for β-catenin destined to enter the nucleus, and that this accumulation of β-catenin levels in the ERC could effectively affects β-catenin substrate levels available for downstream pathways and being critical for cancer progression (28).

In our study, the dual-colour immunofluorescence staining of PI3KC2β with E-Cadherin revealed a partial co-localization of these proteins in a similar discrete perinuclear region (Fig 4A). This supports the hypothesis that PI3KC2β could be promoting the internalization of E-cadherin, β-catenin and EGFR. PI3KC2β is one of the major producers of PI3P in endosomes (29). Moreover, an
*in vivo* study performed by Alliouachene *et al*. reported that the reduction of PI3P upon PI3K-C2β inactivation has selectively impact on endosomal trafficking (11), specifically in the maturation of the APPL1-positive very early endosomes. Moreover, in this study they also observed that PI3K-C2β inactivation led to an accumulation of the insulin receptors in these early endosomal compartments. Therefore, we think that the overexpression of PI3KC2β, probably though an increase in PI3P production, is strongly increasing the level of internalization of E-cadherin, β-catenin and EGFR inducing their accumulation possibly in the ERC. This notably decreases the presence of these molecules in the plasma membrane and adherent junctions, consequently affecting the stabilization of cell-cell adhesion and probably leading to EMT (Fig 7) (28).

In addition to the observed effect in the cellular localization of EGFR, a decrease in EGFR expression was also reported after PI3KC2β overexpression (Fig 5B). This observation is in agreement with previous studies showing that in A431 cells specific cellular context, EMT is also associated with a coordinated loss of EGFR (30). Consistency with the decrease of EGFR, the phosphorylation status of AKT and ERK, downstream targets of this receptor, was also decreased (Fig 5C). This reduction was less remarkable when compared to β-Actin levels, since a strong increase of total-ERK was also found (Fig 5C), suggesting in addition a possible compensatory mechanism.

Together with the decrease in EGFR expression, a reduced sensitivity to the Erlotinib and Gefitinib was also observed (Fig 5D). This is in agreement with the transition to a mesenchymal-like phenotype that is known to decrease the cellular dependence on EGFR signalling, as alternative growth pathways are activated (31) (Fig 7). This has been confirmed in different clinical trials where NSCLC with high expression of E-cadherin showed a beneficial response to Erlotinib treatment in comparison to E-cadherin-negative patients who have an overall deteriorated condition (32). Carcinoma cell lines expressing epithelial proteins, such as E-cadherin, are sensitive to growth inhibition by erlotinib, whereas those tumour cell lines that had undergone an EMT-like transition are less sensitive to EGFR inhibition (33). Similar patterns have been reported for pancreatic and colorectal tumour cell lines (34).

Additionally, PI3KC2β has also been suggested to participate in the resistance to different chemotherapeutic compounds like tamoxifen, cisplatin, Etoposide and Doxorubicin (18,19,35). Moreover, PI3KC2β expression has also been significantly correlated to resistance towards Erlotinib in glioblastoma pathogenesis which supports our results (36).

In HT29 cells the stable overexpression of PI3KC2β was able to reproduce the most notable changes observed in A431 cells. HT29C2β showed increased mRNA levels of Snail and decreased protein levels of E-Cadherin (Fig 6A-B). A change in migration pattern decreasing cell-to-cell contact was also observed in this cancer cell line (Fig 6C). These results further validate our hypothesis where the overexpression of PI3KC2β is promoting EMT.

To conclude, our study shows a link between the overexpression of PI3KC2β, the regulation of intracellular vesicular trafficking and EMT. Further studies are needed to test PI3KC2β as a drug target to revert this malignant phenotype and prevent metastasis formation. Moreover, there is still a necessity to clarify the signalling pathways specifically regulated by PI3KC2β because the recompilation of previous studies suggest that it could be cell/tissue specific. For example, while in some cancers AKT has been proposed as a downstream target (18,19,37), in others this lipid kinase has been associated with MAPK signalling pathway (13), or an alternative miR-449a/β-catenin/cyclin B1 pathway (12). Therefore, the implication of PI3KC2β in EMT could variate depending on the cellular context, and this makes urgent further studies to understand the specific role of this lipid kinase in human biology and tumorigenesis.

## Acknowledgments

We would like to acknowledge the Microscopy Imaging Center of the University of Bern (MIC). Dr Silvia Crespo was enrolled in the Graduate School for Cellular and Biomedical Sciences (GCB) of the University of Bern during the work presented here, we would like therefore to acknowledge it for the training provided. We would also like to thank Dr. Karolina Blajecka for providing the HT29C2β cells and Prof J. Gertsch for his support during the thesis work. We would also like to acknowledge the financial support of the Berner Stiftung für krebskranke Kinder und Jugendliche as well as Batzebär.

